# Parallel and dynamic attention allocation during natural reading

**DOI:** 10.1101/2025.05.27.656336

**Authors:** Yali Pan, Steven Frisson, Joshua Snell, Kara D Federmeier, Ole Jensen

## Abstract

During natural reading, attention constantly shifts across words, yet how linguistic properties (e.g., lexical frequency) impact the allocation of attention remains unclear. In this study, we co-registered MEG data and eye movements while participants read one-line sentences containing target words of either low or high lexical frequency. Using rapid invisible frequency tagging (RIFT), we simultaneously tracked attention to target and post-target words by flickering them at different frequencies. First, we provide neural evidence that attention was allocated simultaneously to both foveal target and parafoveal post-target words. Second, we found an early parafoveal lexical effect, whereby lower frequency targets demanded more attention prior to fixation, and, additionally, a foveal load effect whereby lower frequency targets reduced the amount of attention allocated to post-targets. Furthermore, flexibility in attention shifts between foveal and parafoveal processing correlated with individual reading speed. These results suggest attention is distributed across multiple words and is flexibly adjusted during reading.

## Introduction

What you are doing now, reading, involves moving your eyes to bring the word of interest into foveal vision, then with focused attention, you crack its linguistic code. While reading feels effortless, it requires seamless interaction of oculomotor control, attention shifts, and linguistic processing^1^. There is no doubt that linguistic information in the text impacts how we allocate our attention; for example, when processing a difficult, low lexical-frequency word, we allocate more attention by fixating it longer, demonstrating the classic word frequency effect^2,3^. However, how a word modulates attention across multiple saccades − before, during, and after fixating on it− remains poorly understood. Investigating the dynamics of attention allocation can also provide key insights for shaping theories of natural reading^4^.

To capture the dynamics of attention allocation during natural reading, it is essential to track not only attention to the fixated word but also to adjacent, yet-to-be-fixated parafoveal words. However, conventional techniques in reading research typically do not directly measure attention allocated to parafoveal vision. Behaviourally, eye tracking informs how long our eyes stay on the fixated word, making it difficult to directly measure the processing of parafoveal words. Instead, parafoveal processing is inferred through two key effects: the preview benefit, where parafoveal preview of a word’s linguistic information shortens the fixation duration when it is subsequently processed foveally^5–14^; and parafoveal-on-foveal (PoF) effects, whereby certain properties of a word in the parafovea impact the fixation duration of the currently fixated word^3,15–19^. Thus, eye tracking based evidence relies on indirect inferences based on how parafoveal characteristics affect either the subsequent fixation or the currently fixated word, rather than providing direct measurements of parafoveal processing. Measurements of neural activity through fixation/event-related-potentials (FRPs/ERPs) have also provided evidence for parafoveal processing by aligning brain activity with fixation/word onsets^20–29^ (for a review see^30^). For instance, the amplitude of the N400 has been shown to be modulated by the predictability of the parafoveal word ^24,25,31^. However, the N400 does not directly measure attention allocation; instead, it reflects the retrieval of a parafoveal word’s representation from long-term memory, with less predictable words eliciting more new semantic activation in long-term memory, thus resulting in larger N400 amplitudes^32^. Despite substantial evidence for parafoveal processing, techniques that allow a direct measurement of the attention allocated to both foveal and parafoveal words have been lacking. With the present study we will tackle this longstanding issue.

A recently developed technique, rapid invisible frequency tagging (RIFT), enables the direct and precise measurement of attention during reading, particularly for words in parafoveal vision. RIFT is derived from steady-state-visually-evoked-potentials elicited by stimuli flickering at high frequencies (>= 60 Hz; visually invisible, to minimize interference with ongoing tasks). Neurons in the visual system respond to the visual flicker by generating a signal at the same tagging frequency band that can be measured with EEG or MEG, a phenomenon termed “frequency tagging”^33–39^. These tagging responses increase with attention, making frequency tagging an ideal technique for measuring attention allocation. In a natural reading task, when a word flickers in a sentence, the attention allocated to this word can be directly tracked by the frequency tagged responses. Our previous studies using RIFT demonstrated that lexical^40^ and semantic^41^ properties of a target word modulate attention allocation even before it is fixated. These effects were observed within 100 ms after fixating the pre-target word, when the target word was still in the parafoveal vision. Furthermore, since tagging responses are yoked to a specific flickering frequency, we can track the allocation of attention to multiple words simultaneously simply by flickering them at different frequencies.

In the current study, we applied RIFT to investigate attentional dynamics during natural reading (Figure 1). Specifically, we aimed to address three questions (Figure 2): 1) Can attention actually be allocated to two words simultaneously? 2) How does linguistic information, such as lexical frequency, modulate attention allocation *before* we fixate on that word? 3) and *while* we fixate on that word? This co-registered eye-tracking and MEG study involved participants (*n* = 42) silently reading 188 one-line sentences. Embedded in each sentence were one or two target words with either high or low lexical frequency (Table 1).

**Figure 1.**
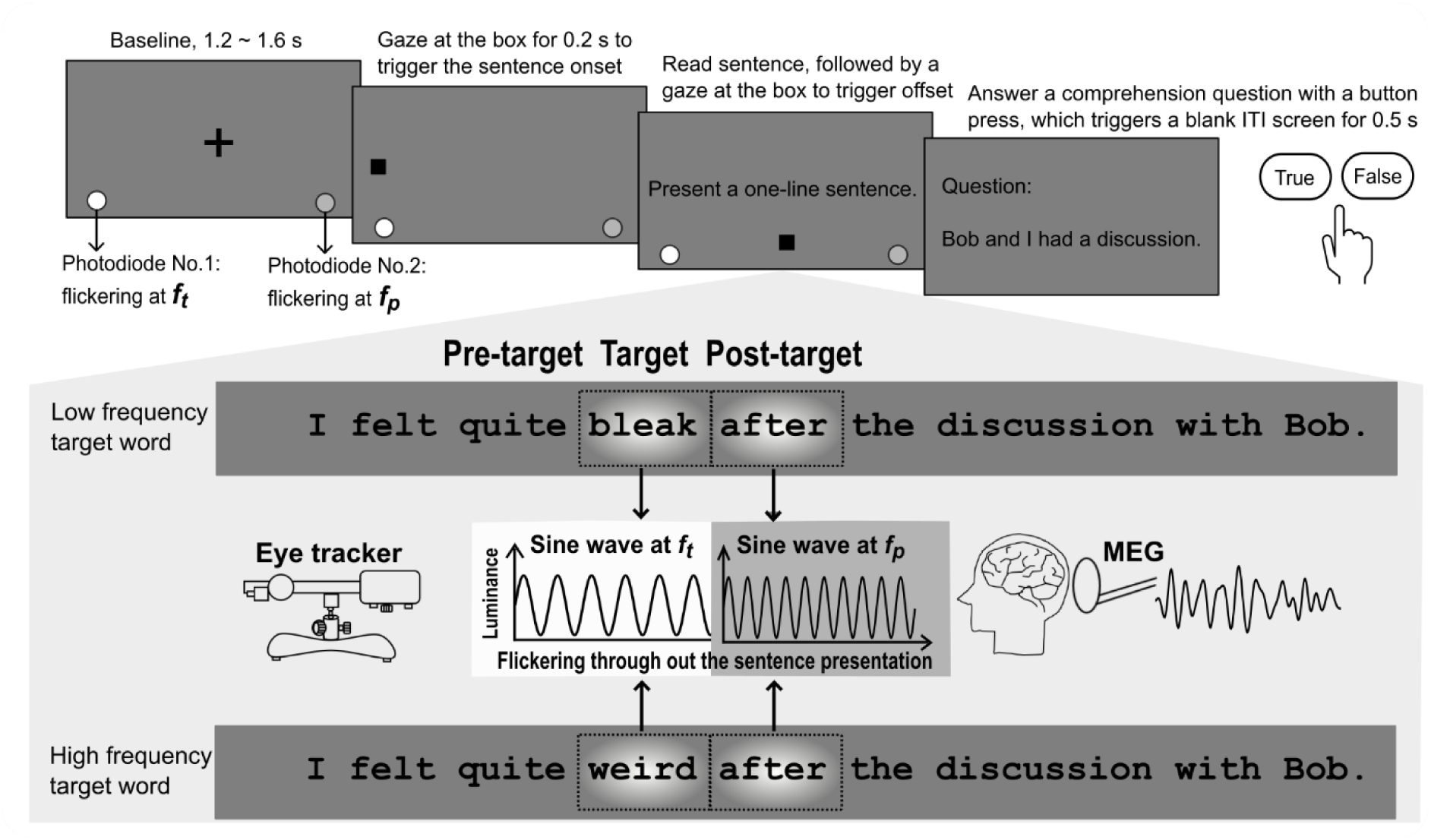
Rapid invisible frequency tagging (RIFT) in a natural reading task. Each trial began with a fixation-cross at the screen centre, followed by a “starting box” on the left. Fixating on the starting box would trigger the sentence onset. Participants (*n* = 42) were instructed to read 188 sentences silently and then gaze at the “ending box” at the screen bottom to trigger the sentence offset. Randomly, 25% of trails included a simple comprehension question requiring a button response. Each sentence contained one or two target words, either of low or high lexical frequency. The target words were unpredictable from the prior context and all sentences were plausible. RIFT was applied by continuously flickering rectangle patches underneath the target and post-target words at frequencies *f_t_* and *f_p_* respectively (60 and 65 Hz sine waves, balanced across participants). A Gaussian mask was applied over the patch to smooth the sharp luminance changes around the edges to reduce their visibility across saccades. Two discs were positioned at the bottom corners of the screen, with their luminance oscillating at sine waves of *f_t_* and *f_p_* separately throughout the entire trial. These discs were covered by two photodiodes, and, during sentence presentation, their luminance changes mirrored those of the patches beneath the flickering words. Eye tracker and MEG data were acquired simultaneously. ITI, inter-trial interval.

**Figure 2.**
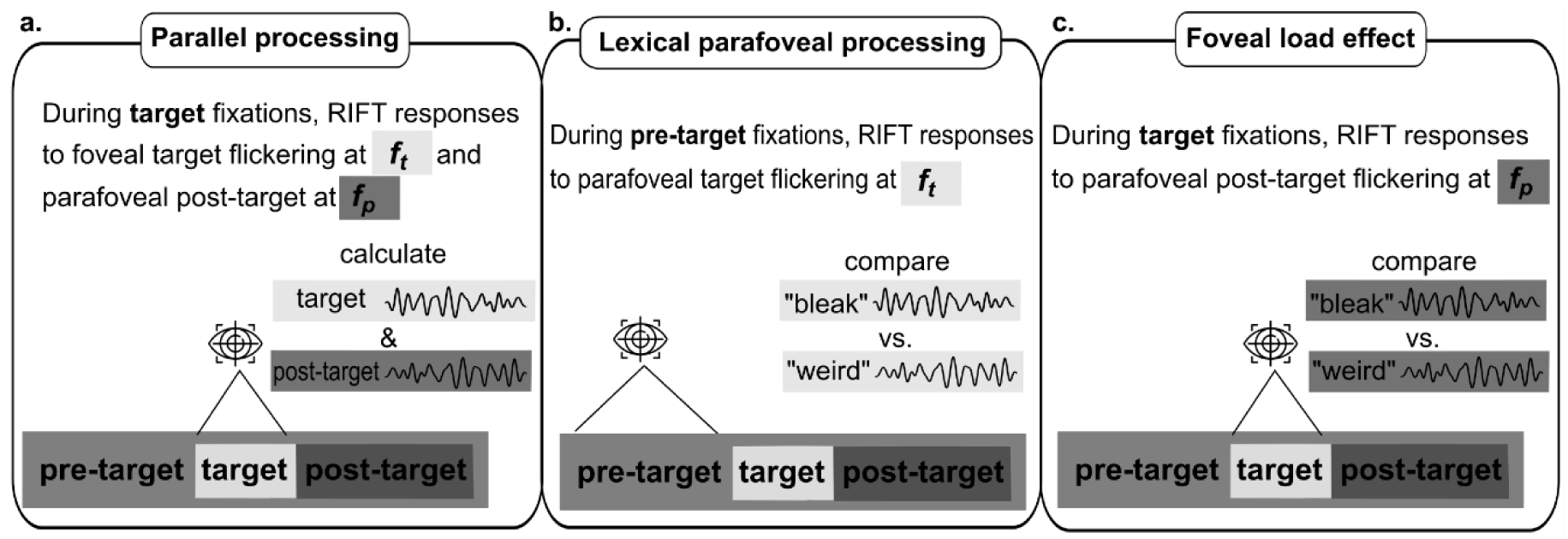
The research questions. **(a)**. When fixating on the target word, the foveal target word was frequency-tagged at *f_t_* and the parafoveal post-target word was frequency-tagged at *f_p_*. By measuring tagging responses associated with *f_t_* and *f_p_*, we could quantify the attention allocated to foveal and parafoveal words, which was used to investigate the parallel processing of multiple words in reading. (**b)**. When fixating on the pre-target word, tagging responses at *f_t_* measured the parafoveal processing of the target word. By comparing the conditional difference, we could probe for lexical parafoveal processing. (**c)**. When fixating on the target word, tagging responses at *f_p_* indicate the amount of attention allocated to the parafoveal post-target word. Conditional contrast of these tagging responses indicates the foveal load effect.

**Table 1.** Word characteristics of pre-target, target, and post-target words.

|  | Pre-target | Target |  | Post-target |
| --- | --- | --- | --- | --- |
| Word frequency | 513.4 (1474.9) | Low | High | 857.5 (2300.4) |
|  |  | 5.5 (4.6) | 91.2 (124.3) |  |
| Word length | 5.7 (1.2) | Low | High | 6.3 (1.8) |
|  |  | 5.7 (0.7) | 5.7 (0.7) |  |

Note. All values here are mean values, with standard deviations in parentheses. Low (<10) and high lexical frequency (>30) target words are reported in terms of the written CELEX frequency per million<sup>42</sup>. Word length refers to the number of letters in a word. The word length of pre-target words ranged from 4 to 8 letters, target words from 5 to 8 letters, and post-target words from 2 to 10 letters. Each low and high-lexical frequency target word pair shared the same pre- and post-target words, balanced across participants.

The target words were unpredictable from the prior sentence context, and all sentences were plausible (see Stimuli and Behavioural pre-tests in Methods). We frequency-tagged both the target and post-target words by simultaneously flickering their respective underlying patches at *f_t_* and *f_p_* (60 and 65 Hz sinusoids, balanced across participants; Figure 1). We also applied Gaussian masks over the flickering patches to minimize their visibility across saccades. Since the flickering frequencies are near or above the critical flicker fusion frequency, participants perceived the flickers as grey, the same colour as the background, rendering them invisible.

RIFT responses provide a direct and precise measure of attention allocated to the target and post-target words throughout the reading process. We hypothesized that: 1) attention would be allocated simultaneously to the target and post-target words, evidenced by significantly-above-baseline RIFT responses at *f_t_* and *f_p_* during the target fixation intervals. 2) The lexical parafoveal effect would result in stronger RIFT responses at *f_t_* when previewing low frequency versus high frequency target words, replicating findings from our previous study^40^. 3) The (word-frequency-based) foveal load effect would manifest as weaker RIFT responses at *f_p_* during fixation on low frequency target words, as greater attention retention in the foveal vision would reduce parafoveal attention available for processing post-target words.

## Results

### No eye movement evidence for parallel word processing in reading

We first examined eye movement measures to test whether we replicated classic effects in the literature — namely, no evidence for lexical frequency effects on eye movement behaviours driven by words in parafoveal vision^43^, but, once those words are fixated, shorter first fixations to higher frequency words. We analysed first fixation durations (i.e., the time spent on a word upon the first landing), an early eye movement measure. No lexical frequency effect was observed for pre-target words (*t*_(41)_ = 0.575, *p* = 0.569; Figure 3a), indicating the lexical frequency of the target word did not affect fixation durations on the pre-target word, providing no evidence for lexical parafoveal processing. As expected, we observed the classic word frequency effect, as evidenced by a significant lexical frequency effect on the first fixation durations of target words, with higher frequency target words associated with shorter durations (*t*_(41)_ = 5.042, *p* = 9.801×10^-6^, two-tailed pairwise *t*-test; Figure 3b). Interestingly, this lexical frequency effect extended to post-target words (*t*_(41)_ = 2.146, *p* = 0.038; Figure 3c), indicating “spill over” of the frequency effect from the target words.

**Figure 3.**
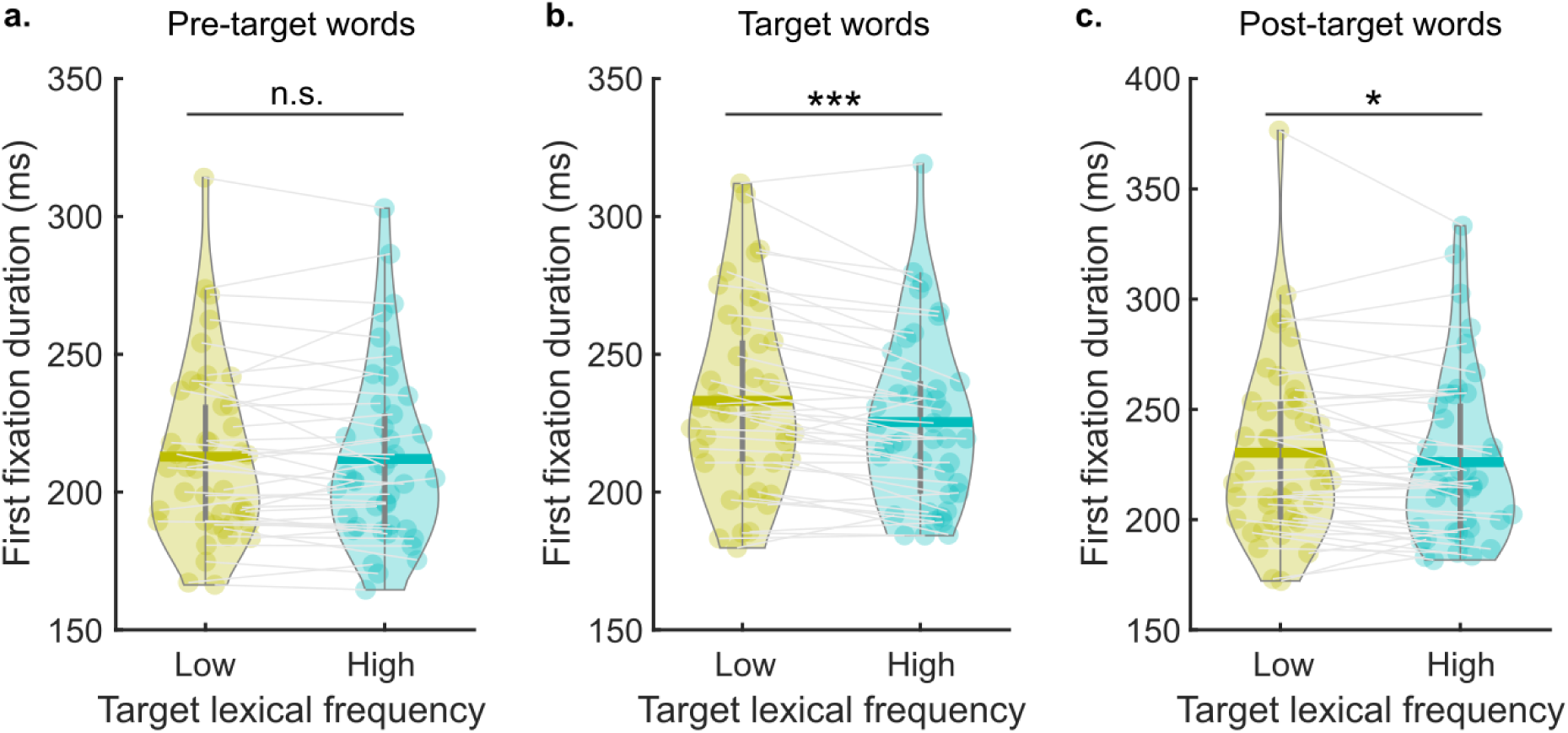
Eye movement results. The first fixation durations of pre-target (**a**), target (**b**), and post-target words (**c**) for conditions of low and high-lexical frequency target words. Each dot denotes the mean of one participant, and the bold horizontal line indicates the overall mean fixation duration over all participants (*n* = 42). Pairwise two-sided *t*-tests were conducted separately for first fixation durations of pre-target, target, and post-target words. \*\*\**p* < 0.001; \**p* < 0.05; n.s., not statistically significant.

### Parallel attention allocation in reading measured by RIFT

MEG data were segmented into a 1 s epoch centred around the first fixation onset of the target words. Coherence between the tagging signal (recorded by a photodiode) and MEG sensors was calculated to measure RIFT responses. Sensors were identified as RIFT response sensors if coherence at the tagging frequency was significantly stronger during target fixation intervals (target word was flickering) compared with baseline intervals (no flickering). This sensor selection procedure included all the trials during each fixation type, irrespective of the experimental conditions (see Methods). Sensor selection was conducted separately for the frequencies of *f_t_* and *f_p_*to ensure precise identification of sensors responding to the foveal and parafoveal tagging (for RIFT sensor topographies see Supplementary figure 1; for *f_t_*, 5.2 ± 5.7 sensors per participant (mean ± STD), range from 1 to 26 sensors; for *f_p_*, 6.3 ± 5.6 sensors, range from 1 to 27 sensors). Further coherence analysis was based on participants with RIFT sensors for both the foveal and parafoveal tagging (29 out of 42 participants).

Target words were frequency tagged at *f_t_* in foveal vision, while post-target words were frequency tagged at *f_p_* in parafoveal vision. Therefore, tagging responses at *f_t_* indicated foveal attention, while tagging responses at *f_p_* indicated parafoveal attention. Using the corresponding RIFT sensors, we calculated coherence as a function of time for the frequencies *f_t_* and *f_p_* (Figure 4). Additionally, we calculated the coherence during the baseline intervals before the sentence presentation to estimate the noise level of coherence. The tagging signal was from the flickering disks on the corners that were covered by photodiodes. We then averaged the coherence values over the 0.2 s target fixations for foveal attention and parafoveal attention, as well as the coherence during the baseline intervals.

**Figure 4.**
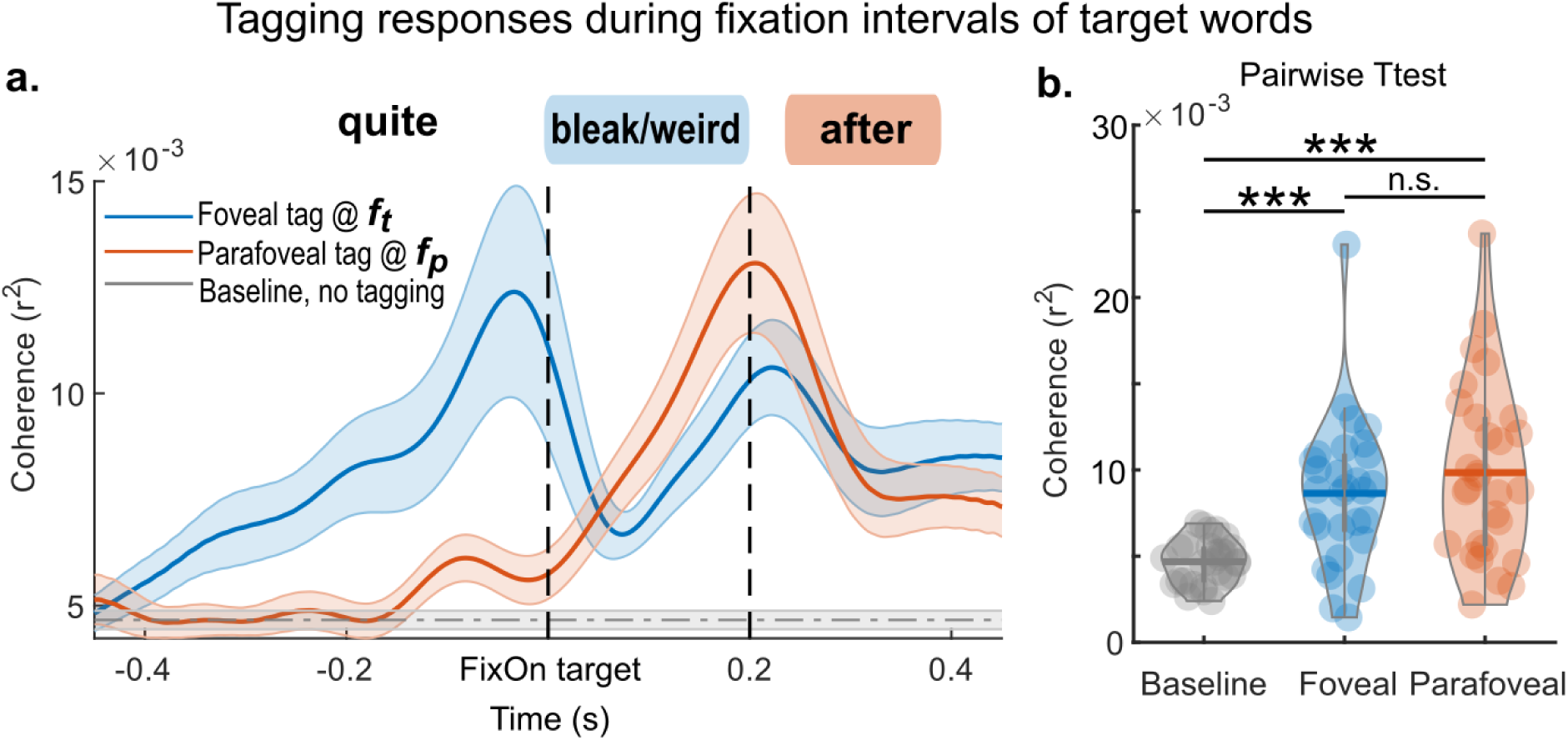
Coherence curves measuring the parallel attention allocated to foveal and parafoveal words. **(a).** Target words were frequency tagged at *f_t_* and post-target words were frequency tagged at *f_p_* (zero time point denotes the first fixation onset of target words). Therefore, RIFT measures (coherence) at *f_t_* (in blue) over all trials measured the foveal attention to the target word, while coherence at *f_p_* (in orange) measured the parafoveal attention to the post-target word. The noise level of the coherence estimation was assessed by calculating the coherence (in grey) during baseline intervals, when only the photodiode discs were flickering, and no flickering words were presented on the screen. Shaded areas around the curve show the standard error around the mean value across participants. (**b)**. Pairwise two-sided *t*-tests were conducted between these three coherence curves (averaged over 0 – 0.2 s). Violin plots display participant-level mean coherence, with bold lines indicating overall averages. \*\*\**p* < 0.001; n.s., not statistically significant.

Dependent sample *t*-tests (pairwise, two-sided) revealed significantly stronger coherence during both foveal (*t*_(28)_ = 4.544, *p* = 9.636×10^-5^) and parafoveal processing (*t*_(28)_ = 5.198, *p* = 1.617×10^-5^) compared with the baseline. Additionally, no significant difference was found between foveal and parafoveal coherence during the target fixation intervals (*t*_(28)_ = 1.415, *p* = 0.168), meaning that the amount of attention allocation to foveal and parafoveal words was comparable. We also questioned whether the parallel allocation of attention as reflected in Figure 4 may have been caused by exclusive coherence at *f_t_* on one half of trials and exclusive coherence at *f_p_* on the other half of trials (which would mean that attention was actually only at one word at any given timepoint). In the Supplementary Notes we provide the single-trial power analysis that rules out this scenario.

In addition to coherence curves at *f_t_* and *f_p_* over all trials during target fixation intervals (Figure 4), we also estimated coherence curves separately by condition (Supplementary figure 2) for pre-target and post-target fixation intervals. For the corresponding RIFT response sensors used in this coherence analysis, see Supplementary figure 1.

### Neural evidence for lexical parafoveal processing in natural reading

Eye movement data showed no evidence of a parafoveal-on-foveal (PoF) lexical effect (Figure 3). The neural data measured as RIFT responses, however, revealed significant differences (Figure 5, column **a**). First, RIFT response sensors were selected based on a stronger coherence at *f_t_* during pre-target fixation intervals (using all trials) compared with the baseline intervals (for RIFT sensors topography see Figure 5 row **i**). Then, for participants with RIFT sensors (30 out of 42 participants), the time-resolved coherence spectra were calculated for the low and high lexical target conditions (Figure 5, row **iii** and **iv**). Afterwards, coherence spectra were averaged over all sensors-of-interest for each RIFT-response-participant. To compare the coherence differences between these two experimental conditions, a group-level cluster-based permutation test was performed during pre-target fixations (0 − 0.2 s) at the frequency of *f_t_* (± 3Hz) (Figure 5, row **v**). We found a significant positive cluster indicating a stronger coherence during pre-target fixations when previewing a low lexical frequency target word (low – high, *p*_cluster_ = 0.036, pairwise two-sided *t*-test during permutations; see Methods for details).

**Figure 5:**
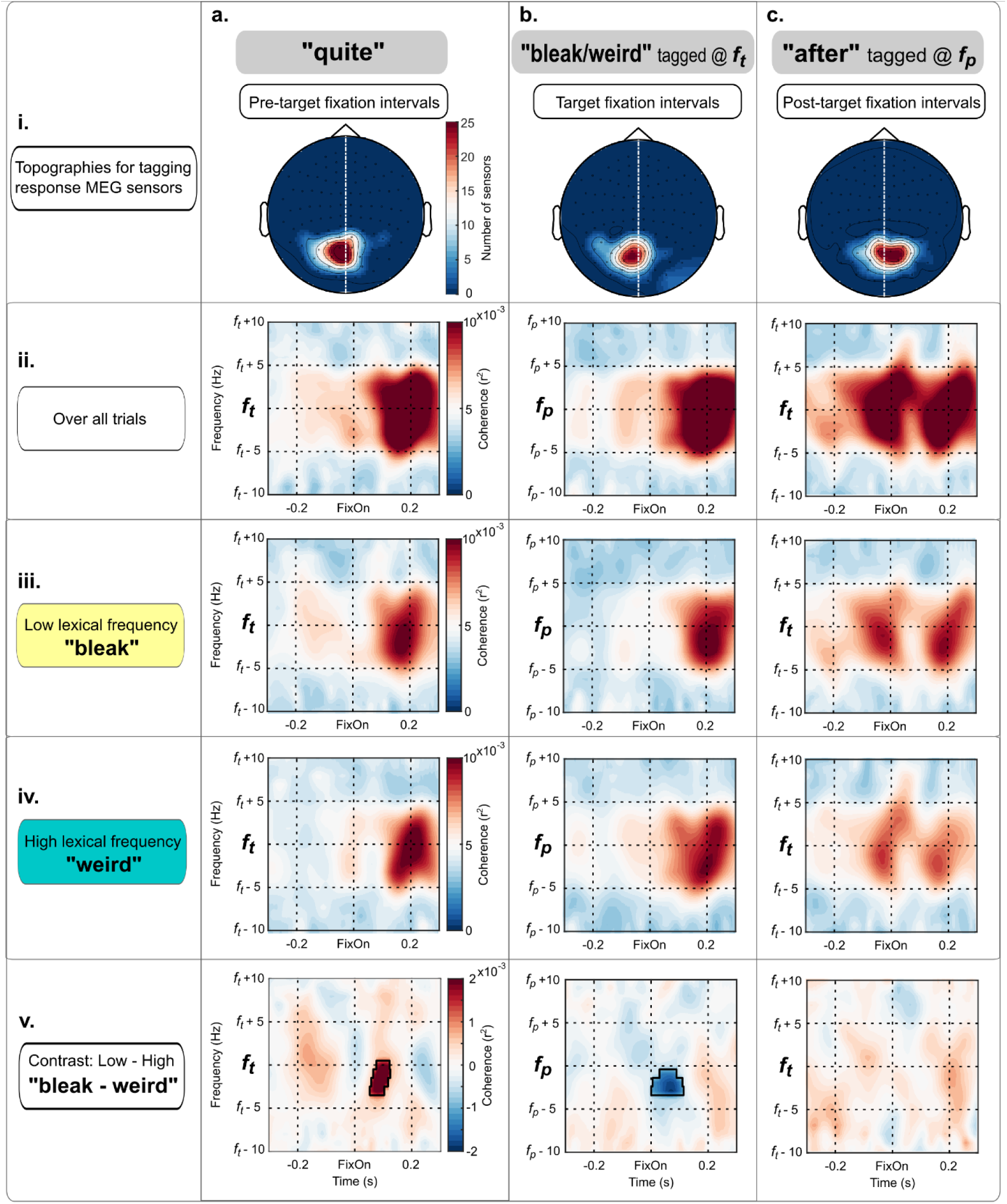
Time-resolved coherence spectra during fixation intervals. First, we selected RIFT response sensors separately for pre-target, target, and post-target fixation intervals by calculating the coherence at their corresponding frequency of interest (*f_t_* for pre-target, *f_p_* for target, and *f_t_* for post-target; columns **a**, **b**, **c**). Topographies for RIFT response sensors summarized over all participants are shown in row **i**. Then, using these RIFT response sensors, time-resolved coherence spectra were estimated for all trials (row **ii**) and for low and high lexical frequency conditions (row **iii**, **iv**). Finally, cluster-based permutation tests were performed to test the conditional difference (row **v**). Significant clusters are highlighted in the coherence spectra by masking non-significant areas (multiply coherence values by 0.4).

### Foveal load determined attention allocated to parafoveal words

We then asked whether the lexical frequency of the foveal word determined how much attention was allocated to parafoveal words. During the fixation intervals of the target word, the post-target word was frequency tagged at *f_p_* in the parafovea (Figure 5, column **b**). First, RIFT sensors were selected based on stronger coherence at *f_p_* during target fixation intervals compared with baseline intervals (for RIFT sensors topography see Figure 5 row **i)**. Then, for the RIFT response participants (33 out of 42 participants), an averaged coherence spectrum over all RIFT sensors was calculated. A group-level cluster-based permutation test during target fixations (0 – 0.2 s) at *f_p_* (± 3Hz) revealed a significant negative cluster (low – high, *p*_cluster_ = 0.025, Figure 5 row **v**). This result demonstrated the foveal load effect: when readers fixated on low lexical frequency words in the foveal vision, less attention was allocated to the parafoveal vision, resulting in weaker tagging responses.

To explore whether attention would regress back covertly to the more difficult, low frequency words, we also did a conditional contrast during the post-target fixation intervals (Figure 5, column **c**). We first selected the RIFT sensors-of-interest during the post-target fixation intervals when the target word was tagged at *f_t_*. Then a cluster-based permutation test was performed, revealing no significant positive or negative clusters (low – high, *p*_cluster_ = 0.290 for the positive clusters; no negative clusters were found), indicating that covert attention did not shift further back when the just-processed word was of low frequency.

### Correlation between attention shifts and reading speed

To examine the behavioural significance of the dynamics of attention shifts, we correlated attentional flexibility with reading speed. Here, flexibility was defined as the difference between the lexical parafoveal and the foveal load effects. The lexical parafoveal effect was quantified as the averaged coherence difference between conditions (low – high parafoveal lexicality) within the significant clusters during pre-target fixation intervals (Figure 5, column **a** row **v**). Similarly, the foveal load effect was quantified as the averaged coherence difference within the significant clusters during target fixation intervals (Figure 5, column **b** row **v**).

Individual reading speed was estimated by averaging the first fixation durations over all words in the sentences, i.e., a lower value indicated a faster reading speed (27 out of 42 participants who responded to both the target and post-target flickers). A Pearson’s correlation analysis revealed a significant negative correlation between the flexibility of attention shift and reading speed (*r* = -0.468, *p* = 0.014, Figure 6).

**Figure 6:**
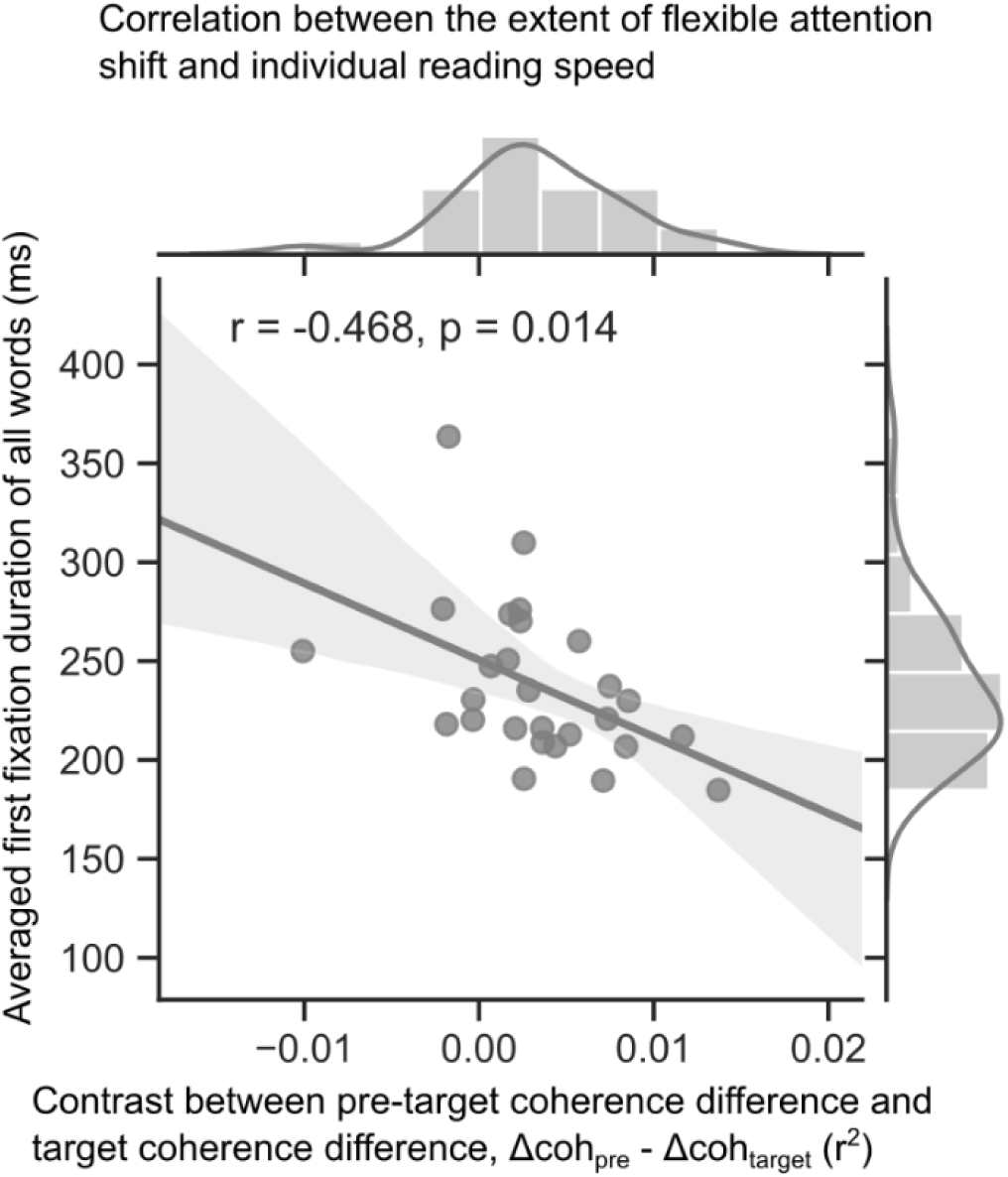
Flexible allocation of attention correlates with reading speed. Flexibility in shifting attention was quantified by contrasting the lexical parafoveal effect with the foveal load effect. The lexical parafoveal effect was derived from the coherence difference averaged over the significant cluster during pre-target fixation intervals (Figure 5, column **a** row **v**), while the foveal load effect was derived from the coherence difference averaged over the significant cluster during target fixation intervals (Figure 5, column **b** row **v**). Individual reading speed was estimated by averaging the first fixation durations over all words in the sentences. We found a significant correlation between flexibility of attentional shifts and reading speed (Pearson’s *r* = -0.468, *p* = 0.014).

## Discussion

How is attention distributed during reading? Here we applied rapid invisible frequency tagging (RIFT) in a natural reading task, simultaneously flickering both the target and post-target words as sinusoids at frequency *f_t_* and *f_p_* (Figure 1) while recording MEG and eye-tracking data. By calculating the RIFT responses, we directly and precisely tracked attention allocated to multiple words (i.e., the target and post-target words) across multiple saccades (i.e., before, during, and after target fixation). First, when fixating on a target word while pre-processing the post-target word in the parafovea, we observed significant RIFT responses to both words (Figure 4), indicating attention being allocated to two words simultaneously.

Turning to the effects of lexical frequency, before fixating on a low frequency target word, we observed stronger RIFT responses at *f_t_*, indicating increased parafoveal attention (Figure 5, column **a**). However, when fixating on this low frequency target word, less attention was allocated to the parafovea, as evidenced by weaker RIFT responses at *f_p_* (Figure 5, column **b**). This flexibility in shifting attention between parafoveal and foveal vision was positively correlated with reading speed, with a higher degree of flexibility linked to faster reading speeds (Figure 6).

First, we provide direct electrophysiological evidence that attention can be allocated to two words simultaneously (Figure 4). Whether attention can be split between foveal and parafoveal words lies at the heart of the debate between serial and parallel processing models. Serial models^44–48^, such as E-Z Reader, assume that attention moves across words in a serial fashion, with only one word under the spotlight of attention at any given time. However, by simultaneously tracking attention to both the foveal and parafoveal words, we found that the attention spotlight can subsume at least two words during a single fixation. Importantly, the simultaneous high tagging responses for foveal and parafoveal words cannot be attributed to a trial-level confound in which attention is focused on the fovea in some trials and on the parafovea in others (see Supplementary Notes and Supplementary figure 3 and 4 for details). The absence of a time shift between the attentional distribution of the two words, as indicated by the overlapping tagging responses curves in Figure 4, suggests that attention can be allocated simultaneously to foveal and parafoveal words, rather than simply shifting to the parafoveal word while the eye remain on the foveal one (Figure 4). When considered alongside the lexical parafoveal effect shown in Figure 5 panel **a** (discussed further in the paragraph below), our results indicate that the amount of attention allocated to the parafoveal word is sufficient for decoding the lexical information. These findings thus point to the recognition of two words during a single fixation, providing compelling neural evidence in support of parallel processing models^49–54^. In the future, theoretical frameworks of natural reading should give greater weight to the flexible, distributed nature of attention and its capacity to support simultaneous processing across multiple words.

If the neural data evidence parallel word processing, why does the oculomotor data not follow suit? In our RIFT-based neural data, stronger RIFT responses were observed for low frequency target words during parafoveal processing (Figure 5, column **a**); in contrast, lexical information from parafoveal words did not influence the fixation durations on the foveal words (Figure 3**a**). This pattern, also observed in our previous study^40^, suggests a dissociation between attention (measured by RIFT responses) and eye movements (measured by fixation durations) during natural reading. That is, attention acts “ahead of” the eyes to pre-process the parafoveal word, but this parafoveal processing need not be reflected in changes in fixation patterns. It is possible that the parafoveal information is either too subtle or too slow to affect saccade programming before the point-of-no-return. Alternatively, it may be that the attentional allocation to the parafoveal word is sufficient for the extraction of key information, such that additional fixation time is not needed. In either case, eye movement patterns do not need to be adjusted word by word, which may serve to optimize cognitive resources for processing words efficiently during reading. This is supported by our correlation results: readers who can flexibly shift attention, without slowing down the eyes (as reflected in the absence of a lexical parafoveal effect in the eye tracking data), tend to process words more quickly (Figure 6). This observation aligns with Yang and McConkie’s model, which proposes that most eye movements are not directly controlled by cognitive processes^55–57^. Furthermore, this notion underpins the principle of autonomous saccade generation in the SWIFT model^50^. In sum, our findings highlight that eye movements do not inform the serial-versus-parallel debate.

The RIFT responses also revealed a foveal load effect: foveal processing of a low compared to high frequency target word reduced attention allocation to parafoveal words (Figure 5, column **b**). This reduction in parafoveal processing due to the increased demands of foveal processing has been repeatedly demonstrated in eye movement studies^58–61^ and event-related-potentials studies^62,63^. In our study, we provide neural evidence for this effect and specify its precise timing. The reduction in parafoveal attention, reflected by reduced RIFT responses, occurred immediately after the fixation onset on the low frequency target word. This immediate modulation of parafoveal attention does not align with the timeline predicted by serial processing models, which assume that attention shifts to the parafovea only after some level of foveal processing has been completed^44–48^. Instead, this rapid modulation of attention suggests that lexical information of the target word was already extracted before fixating on it, as supported by the lexical parafoveal effect discussed earlier. Our previous study even found a modulation of saccade timing based on parafoveal lexical information^64^.

Combining the lexical parafoveal effect and the foveal load effect, we observed a dynamic and flexible shift of attention during natural reading. When the parafoveal word is more difficult to process, more attention is allocated to it; however, while fixating on this difficult word, more attention is withheld to prioritize foveal processing. This flexibility in shifting attention was correlated with reading speed (Figure 6). This correlation also suggests that fast readers do not necessarily always allocate more attention to pre-process parafoveal words; they also retain attention foveally when the currently fixated word is more demanding. Therefore, flexibility, rather than constant forward allocation, may be key to efficient reading.

In summary, by applying RIFT in a natural reading task, we provide the neural evidence that attention can be split between foveal and parafoveal words in a single fixation. Furthermore, we observed that lexical frequency dynamically modulates attention allocation: increased parafoveal attention facilitates previewing of difficult words, while foveal attention retention supports further processing of difficult words. This dynamic flexibility in attention shifts underlies efficient reading. Our study contributes to parallel word processing theories and demonstrates the utility of combining eye-tracking with RIFT to obtain comprehensive insights into the neural underpinnings of reading.

## Materials and Methods

### Participants

Forty-two participants (32 females; age: 22.3 ± 3.3 years, mean ± SD) were recruited for the current study. All participants were native English speakers with normal or corrected-to-normal vision (via contact lenses, not glasses). Participants were right-handed and had no history of neurological conditions or language disorders such as dyslexia. Ethical approval for the study was obtained from the University of Birmingham Ethics Committee (Ethics approval number: ERN-18-0226AP27). Informed consent was obtained from all participants following a full explanation of the study. Compensation was provided as either £30 or 2 course credits. The study lasted approximately 2 hours, including preparation and MEG scanning time.

### Stimuli

In the current study we had 188 sentences, consisting of two sentence sets.

#### First sentence set

The first sentence set consisted of 80 sentences, selected from our previous study^40^. Sentences that were embedded with a long (i.e., more than 8 letters) pre-target word or target word were not selected, as prior findings demonstrated that long pre-target and target words reduce parafoveal processing (see Supplementary figure 5 and Supplementary Notes in^40^).

For the selected sentences, pre-target word lengths range from 4 to 8 letters, while target word lengths ranged from 5 to 8 letters. For a given sentence, the combined word length of pre-target and target words was between 9 to 15 letters, ensuring that the target word fell within the perceptual span and could undergo parafoveal processing. The same word length standard was applied in the second sentence set as well.

Each sentence was embedded with one target word, which was never placed within the first or last three words in the sentence. Target words were always nouns and appeared in grammatical structures comprising adjectives, nouns, and verbs in the pre-target, target, and post-target positions, respectively. These target words varied in lexical frequency (low and high) and were matched for word length (for instance, “ *waltz*” and ”*music*”). Each pair of target words was embedded in two different sentence frames. The surrounding pre-target and post-target words were of the same length across the two sentences frames. The target words within each pair were interchangeable between the two sentence frames, yielding two versions of sentences that were balanced across participants. An example pair of target words for version A and B are shown below (note that target words in the example are bolded and in italics for illustration but were not set apart in the actual experiment). Words were presented in equal-spaced Courier New font.

A. Mike thought this difficult ***waltz*** received lots of criticism.

It was obvious that the beautiful ***music*** captured her attention.

B. Mike thought this difficult ***music*** received lots of criticism.

It was obvious that the beautiful ***waltz*** captured her attention.

In total, the first sentence set contained 80 sentences, embedded with 40 pairs of low and high frequency target words.

#### Second sentence set

The second sentence set included all of the 108 sentences from Degno and colleagues (2019)^18^. These sentences were embedded with target words from various word classes, such as nouns, adjectives, adverbs, and verbs. Each sentence was embedded with two target words of the same lexical frequency condition; for instance, “bleak” and “risky” in the example sentence below are both low frequency words. These two target words were paired with another two target words of the same word length but from the opposite lexical frequency condition; for instance, “weird” and “nasty” are both high frequency words. These two pairs of target words were interchangeable within the same sentence frame, resulting in two versions per sentence (A and B), balanced across participants; see example below (again, targets are in bold and italic in the example only for illustration). Words were presented in equal-spaced Courier New font.

I felt quite **bleak** after discussing that really **risky** subject with Paul.

I felt quite **weird** after discussing that really **nasty** subject with Paul.

In sum, the second sentence set contained 108 sentences, embedded with 108 pairs of low and high frequency target words. Putting the first and second sentence sets together, we had 188 sentences, with 148 pairs of target words, evenly distributed across low and high lexical frequency conditions. Participants read either sentence version A or version B for both sentence sets.

### Behavioural pre-tests

All sentences were plausible and all target words were unpredictable, as established by prior behavioural norming^18,40^, which we summarize here. For detailed information on the first sentence set, please refer to Pan et. Al.^40^; for the second sentence set, please refer to Degno et. Al.^18^.

Plausibility of the sentences was rated on a 7-point scale, with scores ranging from 1 (least plausible) to 7 (most plausible). The experimental sentences were designed to be highly plausible; therefore, filler sentences with low and medium plausibility were included in the norming to occupy the full range of the scale. All experimental sentences achieved plausibility ratings of 5 and above, confirming that the sentences were plausible.

Predictability of the target words were assessed using a cloze test. Participants read sentence fragments consisting of the experimental materials up to, but not including the target words. They were then asked to write down the first word that came to mind that could continue the sentence fragment (without needing to complete the entire sentence). Predictability of a given word was estimated as the percentage of participants who wrote down exactly this word during the cloze test. The predictability of all target words was below 10%, indicating that the target words could not be predicted based on the prior sentential context. Additionally, the highest predictability of any word at the target word location was below 25%, indicating that the sentential contexts at that point were low constraint.

### Experimental procedure

First, participants were required to remove all metal items on the face and body and change into scrubs. They then read and signed the consent form of this study and reviewed the instructions for the reading task. Following preparations for MEG data acquisition (details below), participants were seated in a dimly lit magnetically shielded room (MSR). The MEG gantry was set at a 60° upright angle, fully covering the participant’s head.

All visual stimuli were projected from the stimulus computer, located outside the MSR, onto the participant screen, located inside the MSR, using a PROPixx DLP LED projector (Vpixx Technologies Inc., Canada; for details see Projection of the sentence stimuli). The experimental stimuli presentation scripts were programmed in Psychophysics Toolbox -3 ^65^. The background colour of the screen was middle-grey (RGB [128 128 128]), and words were displayed in black (RGB [0 0 0]). The words were drawn in an equal-spaced Courier New font, at font size 20, with the font style of bold. In the present set up, each letter and space occupied 0.43° visual angle, with the entire sentence spanning no more than 37° of visual angle in the horizontal direction. Sentences were presented as a single line, vertically centred on the screen, starting 2° from the left edge of the screen (Figure 1).

Each trial began with a fixation cross displayed at the centre of the middle-grey screen for 1.2 – 1.6 seconds (randomly selected from a uniform distribution). This was followed by a black square (the “starting square”) with a radius of 1° visual angle, placed at the left vertical centre, 2° from the left edge of the screen. A gaze at the starting square for 0.2 seconds triggered the onset of the sentence. The entire sentence was displayed on the screen at once, with its first letter placed at the position of the starting square. An “ending square” (the same size as the starting square but grey, RGB [64 64 64]), was presented below the sentence.

Participants were instructed to gaze at the ending square after reading the sentence. A gaze at the ending square for 0.1seconds triggered the offset of the sentence presentation. The trial ended with a blank middle-grey screen that lasted 0.5 seconds. Randomly, in 25% of trials, a comprehension statement about the content of the immediately preceding sentence appeared. Participants were required to answer “True or “False” by pressing a button. The behavioural performance on this comprehension task was high (94.1% ± 5.4%, mean ± SD), indicating that participants read the sentences attentively.

The experiment was divided into four blocks, each lasting approximately 6 minutes. Participants had at least a one-minute break between blocks and could resume the experiment at their convenience by pressing any button. Participants were instructed to read sentences silently at their own pace. In total, the entire experiment took no longer than 40 minutes. Eye movements and brain activity were recorded simultaneously throughout the session, and eye data were co-registered with MEG data in later analysis.

### Rapid invisible frequency tagging (RIFT)

#### Projection of the sentence stimuli

The refresh rate of the PROPixx projector was 1440 Hz (VPixx Technologies Inc., Canada), while the refresh rate of the stimulus computer monitor was 120 Hz (1920 × 1080 pixels resolution). To achieve the high refresh rate of the PROPixx projector at 1440 Hz, the sentence was displayed repeatedly in four quadrants of the stimulus computer monitor. The projector then interpreted these 12 colour channels (3 RGB channels × 4 quadrants) as 12 individual grayscale frames, which were projected onto the participant screen in rapid succession. Consequently, one frame from the stimulus computer became 12 frames for the PROPixx projector, resulting in displaying sentences at 12 × 120 Hz, which was 1440 Hz.

### Frequency tagging of target and post-target words simultaneously

To frequency-tag a word, a rectangular patch was added underneath it. The side length of the patch was the width of this word plus the spaces on both sides, spanning approximately 3° to 4.5° visual angle. We flickered the patch by changing the luminance of each pixel inside of it from black to white in a sinusoid pattern. Using this method, we tagged both the target and post-target words simultaneously by flickering the patches underneath at two different frequencies. For half of the participants, target words were tagged at 60 Hz, and post-target words at 65 Hz; for the other half, target words were tagged at 65 Hz and post-target words at 60 Hz. These two tagging frequencies were balanced across participants rather than trials to ensure maximum number of trials for an optimal signal-to-noise ratio for detecting the tagging responses. For illustration purposes, the tagging frequency of the target words is denoted as *f_t_*, and that of the post-target words was denoted as *f_p_* (Figure 1). The flickering patches were perceived as middle-grey (matching the screen background) due to their sinusoidal luminance modulation, rendering them invisible to participants. Note that both the target and the post-target words were displayed in black and were not flickering, which were the same as the other words on the screen.

The two rectangular patches underneath the target and post-target words were flickering throughout the entire presentation of the sentence, with participants making continuous saccades across words. To minimize sharp luminance transitions and prevent visibility of patch edges during saccades from the non-tagged to the tagged area, we applied a Gaussian-smoothed transparent mask over the flickering patch. The mask was fully transparent at the centre and gradually became opaque towards the edges. The mask was generated using a two-dimensional Gaussian function (Equation 1):

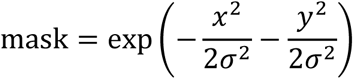

where 𝑥 and 𝑦 are the mesh grid coordinates for the flickering patch, and *σ* is the 𝑥 and 𝑦 spread of the mask with *σ* = 0.02° visual angle.

To record the tagging signal for later coherence analysis, we used two custom-made photodiodes (Aalto NeuroImaging Centre, Finland) to record the luminance changes of the flickering patches. These photodiodes, placed to the bottom-left and bottom-right corners of the participant screen, monitored disks that flickered identically to the flickering patches underneath the target and post-target words. These photodiodes converted luminance/light changes into voltage signals, which were recorded as two external channels in the MEG system, sampled at 1,000 Hz, the same rate as the MEG sensors.

### Data acquisition

#### MEG data

During the whole session of the reading experiment, MEG data were collected using a 306-sensor TRIUX Elekta Neuromag system, which consisted of 204 orthogonal planar gradiometers and 102 magnetometers (Elekta, Finland). To prepare for MEG data acquisition, we first attached four head-position indicator (HPI) coils to each participant’s head — two on the left and right mastoid bone and two on the forehead, with a minimum separation of 3 cm between them. Afterwards, a Polhemus Fastrack electromagnetic digitizer system (Polhemus Inc, USA) was used to digitize the individual head. We first digitized the locations of three bony fiducial points — the nasion, and left and right preauricular points. Then we digitized locations of the four HPI coils. Finally, we digitized the whole head by sampling at least 200 points that covered the whole scalp and were distributed evenly. These points from the digitization procedure were later used in spatially co-registering the MEG head model with individual structural MRI images for the source analysis. During this preparation, we also attached three pairs of electrodes on each participant’s face; one pair was placed above and below the right eye to record the vertical electrooculogram (EOG), another pair was placed 1 cm away from the left and right of the eyes to record the horizontal EOG, and the last pair was placed on the left and right collarbone to record the heart’s electrical activity (i.e., electrocardiogram, EKG or ECG).

After the preparation, participants were seated upright under the MEG gantry at a 60° angle within the magnetically shielded room. The MEG data were sampled at 1,000 Hz, with an online band-pass filtering from 0.1 to 330 Hz to minimize aliasing effects.

### Eye movement data

Eye movements were tracked using an EyeLink 1000 Plus eye-tracker (long-range mount, SR Research Ltd, Canada) throughout the whole MEG session. The eye tracker was placed on a wooden table in front of the participant. The centre of the horizontal bar of the eye tracker was aligned to the middle of the participant screen, and the top of the eye tracker was at the same level as the screen bottom edge. The distance between the eye-tracker camera and the participant’s eyes was 90 cm. We recorded the horizontal and vertical positions as well as the pupil size from the left eye, at a sampling rate of 1,000 Hz. Each block began with a five-point calibration and validation test. The test was accepted if the validation error was below 1° visual angle both horizontally and vertically. During the block, we performed a one-point drift checking test every three trials to ensure a good enough precision of the eye tracking. If the drift check failed or the sentence presentation was unable to be triggered by participant gaze at the starting box, the five-point calibration and validation test was conducted again.

### Eye movement data analysis

We used the EyeLink built-in algorithms to parse eye movement events based on the online detection of saccade onset using the following parameters: the motion threshold as 0.1°, the velocity threshold as 30°/second, and the acceleration threshold as 8,000°/second^2^. This conservative setting is suggested by the EyeLink user manual for reading studies where saccades are prominent, as this parameter setup can prevent false saccade detections and reduce the number of micro-saccades.

Fixation onset events were extracted from the EyeLink output file using custom-made scripts, which were parsed using the parameters described above. A fixation event was assigned to a given word if the averaged x and y positions of this fixation landed in the area of this word and the space to the left of it, because the optimal landing position is slightly left within a word. For further eye movement data analysis, only the fixation that first landed on a given word was selected; i.e., the first fixations. First fixation durations that were shorter than 0.08 second or longer than 1 second were viewed as outliers and discarded from further analysis. For each participant, the averaged first fixation durations for the pre-target, target, and post-target words were obtained separately for the low and high-lexical frequency condition of the target words. Afterwards, dependent sample *t*-tests (pairwise, two-sided) were conducted separately on the first fixation durations of pre-target, target, and post-target words.

### MEG data analysis

The MEG data analysis was conducted using MATLAB R2020a (Mathworks Inc, USA), incorporating the FieldTrip toolbox (version 20200220; Oostenveld et al., 2011), the FLUX MEG analysis pipeline ^67^, and custom-made scripts.

#### Pre-processing

First, malfunctioning sensors were identified through visual inspection, based on excessive high frequency noise compared to other sensors, and then excluded from data analysis (0 to 2 sensors excluded per participant). The MEG data were then band-pass filtered from 0.5 to 100 Hz using phase-preserving, two-pass Butterworth filters. To factor out slow drifts, the data were detrended. In order to remove oculomotor and heartbeat related artefacts, the MEG data were decomposed using an independent component analysis (ICA)^68^. The number of components matched the number of valid MEG sensors (up to 306). Since independent components were ranked by their contribution to explaining the data, we visually inspected only the first 100 components for each participant. We removed only components related to blinks, eye movements, and heartbeat, based on visual inspection of their topographies, power spectrum, and time course (for tutorial see Fieldtrip^66^; we excluded 2 to 5 components per participant).

MEG and eye movement data were co-registered by aligning triggers from the MEG system and the EyeLink device. Afterwards, eye movement events were used to segment the MEG data. MEG data were segmented from -0.5 to 0.5 seconds relative to the onset of the first fixations on the pre-target, target, and post-target words. Segments with extreme fixation durations, i.e., shorter than 0.08 seconds or longer than 1 second, were deemed outliers and excluded from further analysis. Additionally, we segmented 1 second long baseline data, spanning from 0 to 1 second relative to the onset of the fixation cross before the sentence presentation. Through visual inspection of all segments in a condition blind fashion, segments that were contaminated by muscle or movement artefacts were identified and removed from further analysis.

#### Coherence calculation

The brain responses to flickering target and post-target words were quantified separately by calculating coherence between the MEG sensors and the corresponding photodiode channel (the tagging signal). Amplitudes of the photodiode channel were normalized within each segment to standardise photodiode values across all segments and participants. To estimate the time-resolved coherence spectrum for a given frequency of interest, we first filtered the segments using Hamming-tapered Butterworth bandpass filters (4^th^ order, phase preserving, two-pass), with a bandwidth of ± 5 Hz around the centre frequency point. Then, the Hilbert transform was applied to obtain analytic signals of the filtered narrow band data. Afterwards, the analytic signals were used to estimate the coherence at the frequency of interest (Equation 2):

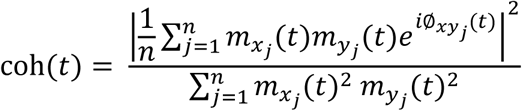

where *n* is the number of segments. For the time point *t* in the segment *j*, 𝑚_𝑥_(𝑡) and 𝑚_𝑦_(𝑡) are the time-varying magnitude of the analytic signals from a MEG sensor (*x*) and the photodiode (*y*) respectively, ∅_𝑥𝑦_(𝑡) is the phase difference as a function of time (for detailed description, please see ^69^.

This coherence analysis was repeated for frequencies of interest from 50 to 75 Hz in a step of 1 Hz to obtain a complete spectrum of time-resolved coherence.

#### Selection for RIFT response sensors

Not all MEG sensors respond to visual flickers, so as the first step of analysing MEG data, RIFT response sensors were selected to enhance analysis sensitivity. This sensor selection procedure was carried out separately for fixation intervals of pre-target, target, and post-target words (for topographies of all three types of sensors see Figure 5 panel i). MEG sensors that showed significantly stronger coherence at *f_t_* during pre-target fixation intervals (when target words were frequency-tagged at *f_t_*) compared with baseline intervals (when nothing was frequency-tagged) were selected as the pre-target RIFT response sensors. Similarly, sensors that showed significantly stronger coherence at *f_p_* during target fixation intervals (when post-target words were frequency-tagged at *f_p_*) compared with baseline intervals were selected as the target RIFT response sensors. Finally, in order to investigate whether attention would regress back covertly to the previous difficult word (i.e., the low frequency target words), we also selected post-target RIFT response sensors, which showed significantly stronger coherence at *f_t_* during post-target fixation intervals (when target words were frequency-tagged at *f_t_*) compared with baseline intervals.

To estimate statistical significance during the RIFT response sensors selection procedure, we used a non-parametric Monte-Carlo method ^70^ to compare the coherence difference between the frequency tagging intervals versus the no-tagging intervals (the baseline segments). The statistical comparison was conducted separately for the three types of frequency tagging intervals — the pre-target, target and post-target fixation intervals. Because previous RIFT studies with a natural reading task only observed robust tagging responses to visual flicker from the visual cortex ^40,41^, this sensor selection procedure was restricted to MEG sensors in the visual cortex (52 planar sensors).

Frequency tagging intervals include all trials, regardless of the lexical conditions of the target words in the experimental design. In the non-parametric statistical analysis here, frequency tagging intervals (the pre-target or target or post-target fixation intervals) and baseline intervals were treated as two conditions. For each combination of the MEG sensor and the photodiode channel, coherence at the frequency of interest was estimated over trials for the frequency tagging and baseline conditions separately. Afterwards, we calculated the z-statistic value for the coherence difference between these two conditions using the following equation (for details please see ^70^) (Equation 3):

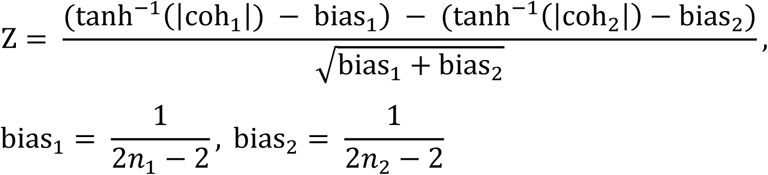

where coh_1_ and coh_2_ denote the coherence value for the frequency tagging and baseline conditions, bias_1_ and bias_2_ is the term used to correct for the bias from trial numbers of the frequency tagging (𝑛_1_) and baseline conditions (𝑛_2_).

After obtaining the z statistic value for the empirical coherence difference (i.e., without shuffling trial labels), we ran a permutation procedure to estimate the statistical significance of this comparison. We randomly shuffled trial labels between the frequency tagging and baseline conditions for 2,000 times. During each permutation, coherence was computed for both conditions using trials with the shuffled labels, then entered into Equation 3 to obtain a z score for the coherence difference. After the 2,000 iterations of permutations were performed, all the z-values were sorted from the minimum value to the maximum value, establishing the null distribution for statistical analysis. Because we had a prior direction of the comparison, i.e., a RIFT response sensor was supposed to have stronger coherence for the frequency tagging condition compared with the baseline condition, the statistical test was one-sided rather than two-sided. If the z-value of the empirical coherence difference exceeding the 99^th^ percentile of the null distribution, this sensor was selected as a RIFT response sensor (right-sided, *p* = .01).

#### Coherence comparison between experimental conditions

The coherence comparison between the two experimental conditions (the low and high-lexical frequency of target words) were only conducted for participants with RIFT response sensors. For each participant, this coherence comparison was performed separately for the fixation intervals of pre-target, target, and post-target words using their corresponding RIFT response sensors. The time-resolved coherence spectrum, derived from Equation 2, were averaged across the RIFT responses sensors, so that each participant had one coherence spectrum for each condition during each frequency tagging interval. Please note that to avoid any bias from unequal trial numbers, we randomly discarded the redundant trials from the condition with more trials, ensuring an equal number of trials per condition were entered the coherence analysis.

To compare the time-resolved coherence between two conditions on the group level, we conducted a cluster-based Monte-Carlo permutation test ^70^. This non-parametric permutation test was performed separately for the three tagging intervals — the fixation intervals of pre-target, target and post-target words. We pre-defined the following frequency window and time window for the permutation test. A frequency window of ± 3 Hz around the frequency of interest was selected because the difference between these two tagging frequencies was 5 Hz; a time window of 0 to 0.2 seconds around the fixation onset of the word of interest was selected because the averaged fixation duration of a given word was around 0.2 seconds.

During each permutation iteration, trial labels between the two experimental conditions were randomly shuffled across participants. This label shuffling could break down the corresponding relation between the coherence values and the conditions, which was essential in creating the null distribution on the group level. Dependent samples *t*-tests were conducted between the coherence values from the two conditions (within the pre-defined frequency and time window, see above) to obtain an array of *t*-values. Then, a significance threshold of 0.05 (two-sided) was applied to cluster the *t*-values. The *t*-values within each cluster were added up to get a summed *t*-value, and then the maximum summed *t*-value was used to create the null distribution (the “maxsum” method of cluster-based permutation test). After 5,000 iterations of permutation, a null distribution of maximum summed *t*-values was created. We then sorted the distribution from the smallest value to the biggest value, and the summed *t*-values at the position of 2.5% and 97.5% were selected as the critical values at the significance level of 0.05 (two-sided). Next, we calculated the dependent samples *t*-test using the empirical data and applied a significance threshold of 0.05 (two-sided) to cluster the *t*-values. By comparing the *t*-values clusters against the critical values from the null distribution, only the significant clusters remained, as outlined in Figure 5 row **v**.

#### Simultaneous coherence analysis during the foveal and parafoveal reading

In order to estimate the dynamics of attention allocated to process the target (frequency tagged at *f_t_*) and post-target words (frequency tagged at *f_p_*) during reading, we calculated tagging responses with respect to *f_t_* and *f_p_* in three time-windows: the fixation intervals of pre-target, target, and the post-target words.

First, using the procedure described in the section of “Selection for the RIFT response sensors”, we selected RIFT response sensors at two tagging frequency bands (*f_t_* and *f_p_*) during three fixation intervals (pre-target, target, and post-target fixations). This sensor selection procedure was conducted over all trials, irrespective of the experimental conditions. Topographies for these six types of RIFT response sensors are provided in Supplementary figure 1. Notably, the topographies of pre-target RIFT sensors at *f_t_*, target RIFT sensors at *f_p_*, and post-target RIFT sensors at *f_t_* were the same as displayed in Figure 5. The selection accounted for frequency-specific interests during each fixation interval to ensure a precise and accurate tagging response detection.

Then, using these selected RIFT sensors, we calculated tagging response curves for each experimental condition following Equation 2. For each type of fixation interval and each condition, we obtained a coherence curve at the frequency of interest (with a bandwidth of 5Hz in the Hilbert filtering). Coherence curves were averaged across RIFT sensors to derive participant-specific curves, which were then averaged across participants. Supplementary figure 2 shows the averaged coherence curves for each fixation interval and each experimental condition. As a no-tagging baseline, we calculated tagging responses during the fixation-cross intervals before the sentence presentation, using post-target RIFT sensors.

Since no flickering occurred during baseline intervals, the choice of RIFT sensors from any fixation intervals did not affect the coherence values, which reflected only chance or noise levels.

During target fixation intervals, target words were frequency tagged at *f_t_* while post-target words were frequency tagged at *f_p_*. Thus, tagging responses from all trials at *f_t_* and *f_p_* were considered as measures for attention allocated to the foveal and parafoveal vision. To assess the significance of tagging responses during foveal and parafoveal processing, we conducted group-level paired *t*-tests, comparing coherence values for target fixation intervals against baseline intervals. The coherence values were averaged over a 0.2 second time window aligned with the target word fixation onset or the baseline presentation onset.

## Data availability

We have deposited the following data in the current study on figshare (https://figshare.com/projects/Attention_in_Reading/250316): the raw MEG data, the epoch data after pre-processing, the raw EyeLink files, the Psychotoolbox data.

## Code availability

The experiment presentation scripts (Psychtoolbox-3), statistics scripts (R), data analysis scripts (MATLAB) are available on GitHub (https://github.com/yalipan666/Attention_in_Reading).

## Acknowledgements

This work was supported by a Leverhulme Early Career Fellow to Y.P. (grant number ECF-2023-626) and a Wellcome Trust Discovery Award to O.J. (grant number 207550).

## Author Contributions

Y.P. devised and designed the study, programmed and conducted the experiment, Y.P. carried out the analyses with assistance from O.J., S.F., J.S., and K.D.F. Y.P. wrote the first draft. Y.P., S.F., J.S., K.D.F., and O.J. edited the manuscript together.

## Supplementary Materials

**Supplementary figure 1.**
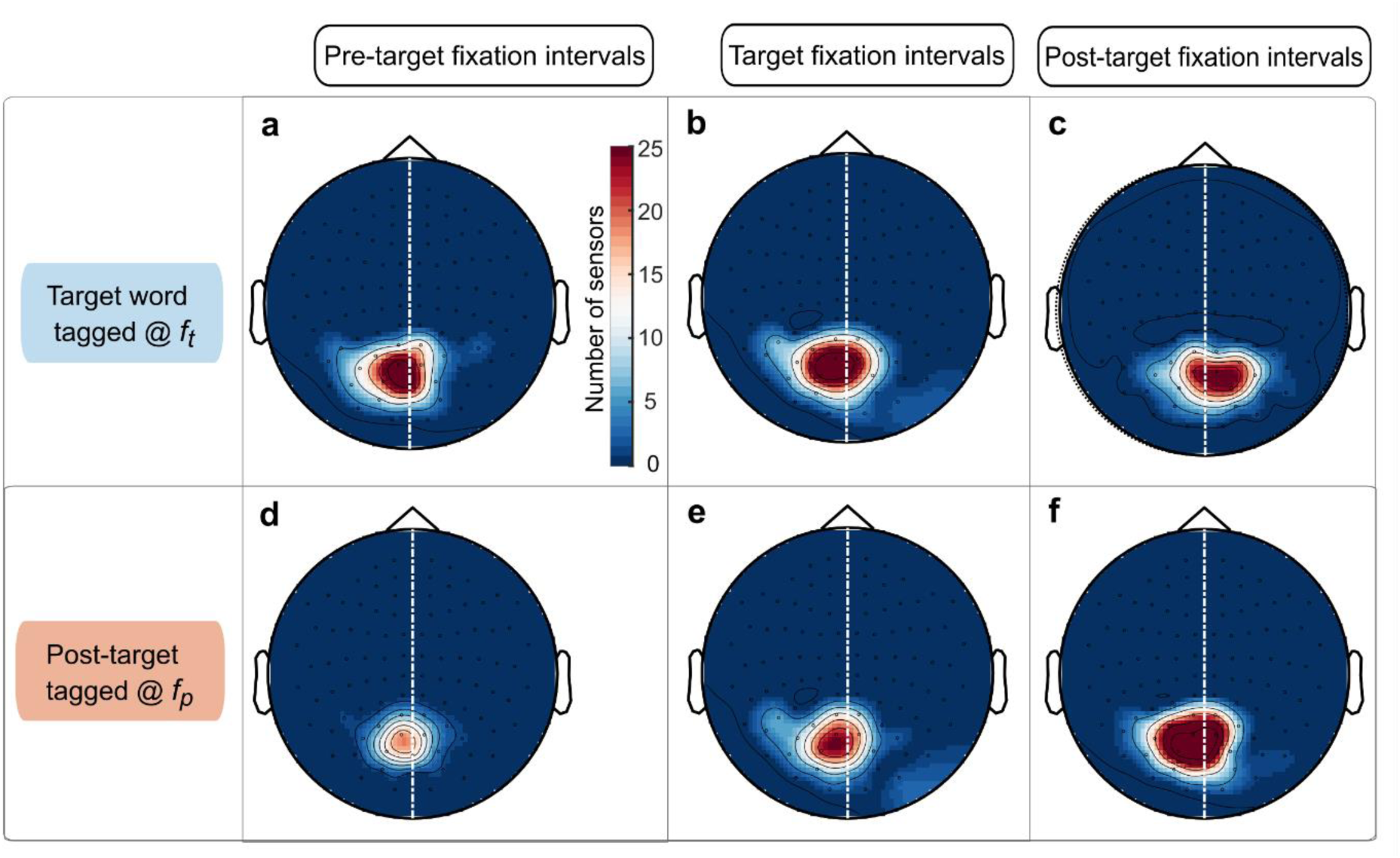
Topographies for rapid invisible frequency tagging (RIFT) response sensors. Using the sensor selection procedure detailed in the section Methods, we identified RIFT response sensors at the two frequency bands of interest (*f_t_* and *f_p_*, the first and second row) during fixation intervals of pre-target, target, and post-target words (the first, second, and third column). In the current experiment, target words in the one-line sentences were frequency tagged at *f_t_*, while post-target words were frequency tagged at *f_p_*. Please note that for this RIFT response sensor selection, we included all trials, regardless of experimental conditions. The topographies summarize the distribution of RIFT sensors over all participants (*n* = 42). Only participants with RIFT sensors were included in further analyses (*n* = 30 in panel **a**, *n* = 32 in panel **b**, *n* = 32 in panel **c**, *n* = 25 in panel **d**, *n* = 33 in panel **e**, *n* = 39 in panel **f**). Panel **a**, **e**, and **c** are the same topographies shown in the Figure 5 row **i**.

**Supplementary figure 2.**
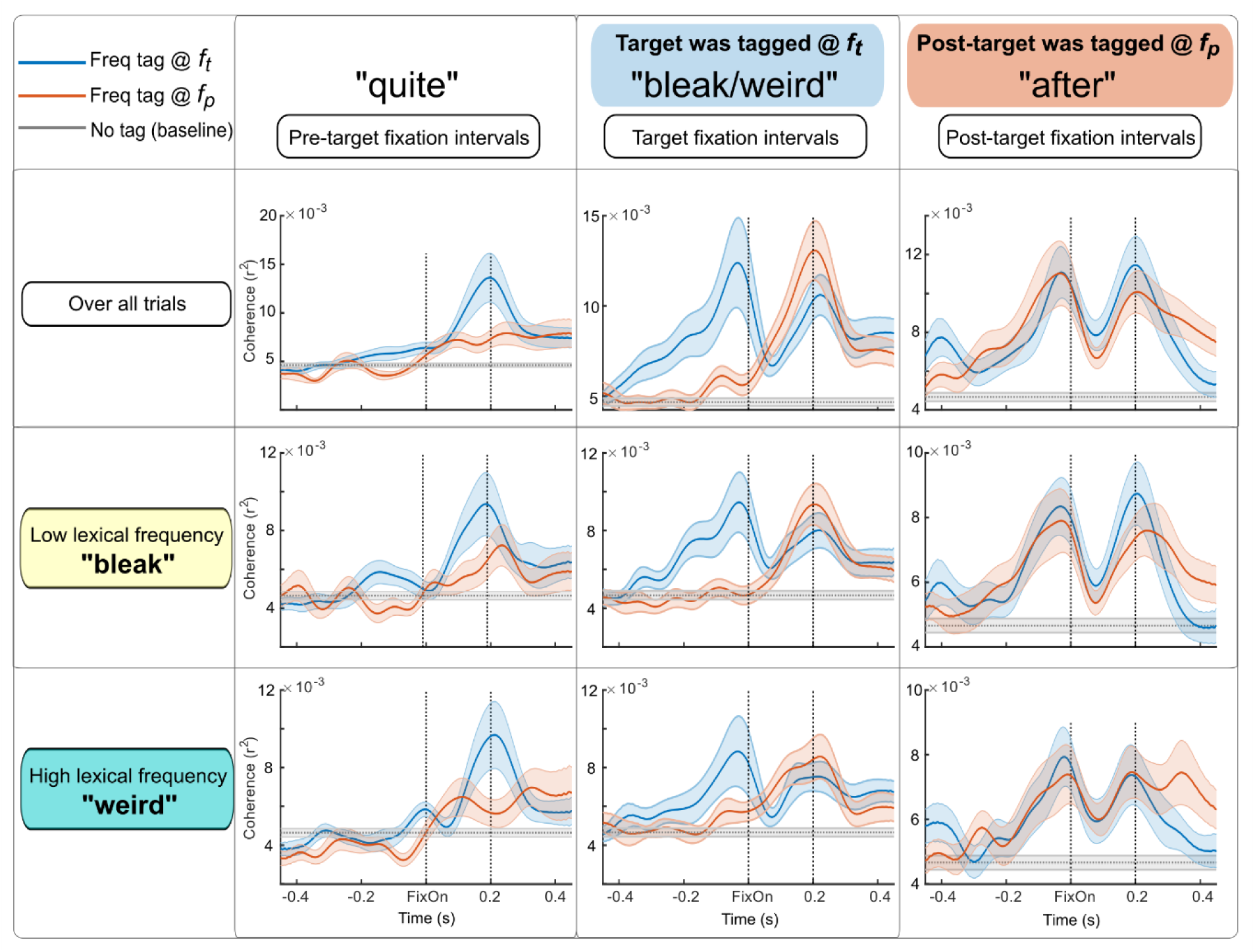
Coherence curves depicting brain responses to RIFT. Using RIFT response sensors (for topographies see Supplementary figure 1), we plotted the brain responses with respect to the tagging frequency of *f_t_* (in blue) and *f_p_* (in orange) during fixation intervals of pre-target, target, and post-target words (zero time-point indicates the first fixation onset to the corresponding word). For each type of the fixation interval, coherence curves were calculated for all conditions, the condition of low lexical frequency, and the condition of high lexical frequency. Additionally, we also calculated coherence curves during baseline intervals, when no flickering occurred, serve as the noise level estimation (in grey). Shaded areas around the curves represent standard errors across participants.

**Supplementary figure 3.**
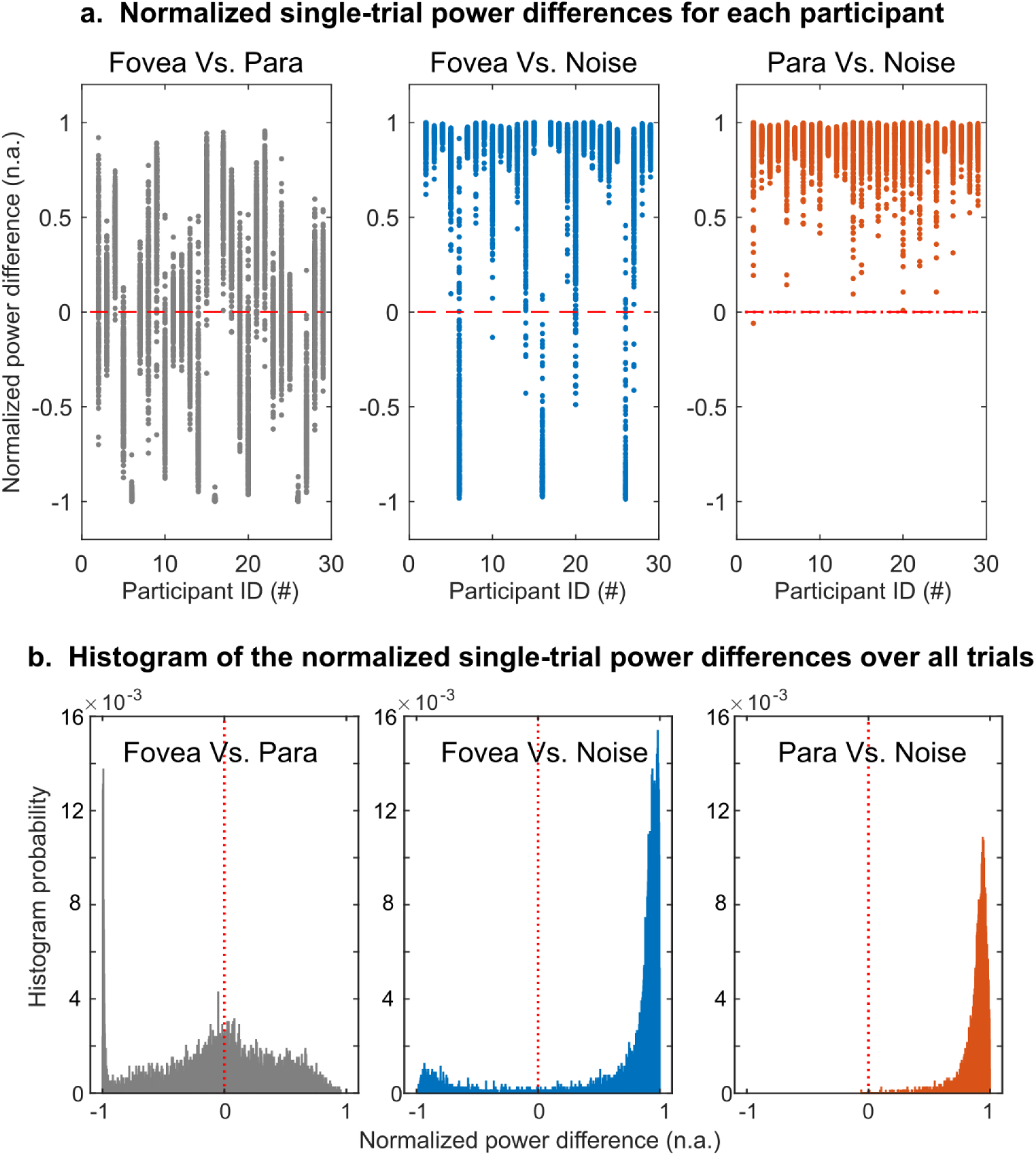
Control analysis using single-trial power confirms parallel attention allocation. To address the possibility that cross-trial coherence may mask trial-level variability in attention allocation, we computed single-trial power at the foveal (*f_t_*) and parafoveal (*f_p_*) tagging frequencies for participants with robust RIFT responses at both frequencies (*n* = 29). The normalised power difference between foveal and parafoveal tagging was calculated for each trial as 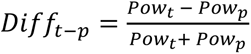, and then averaged across trials and across the time window of 0 – 0.2s (aligned with fixation onset to target words). To confirm that these responses were above noise level, we compared single-trial power at tagging frequencies to power at a non-tagged frequency (i.e., 55 Hz). (**a).** Distributions of the normalized single-trial power differences: between fovea and parafoveal (grey), between fovea and noise (blue), and between parafovea and noise (orange). Each dot indicates power difference of one trial, each column indicates one participant. (**b).** Histograms of these single-trial differences, pooled over all trials.

**Supplementary figure 4.**
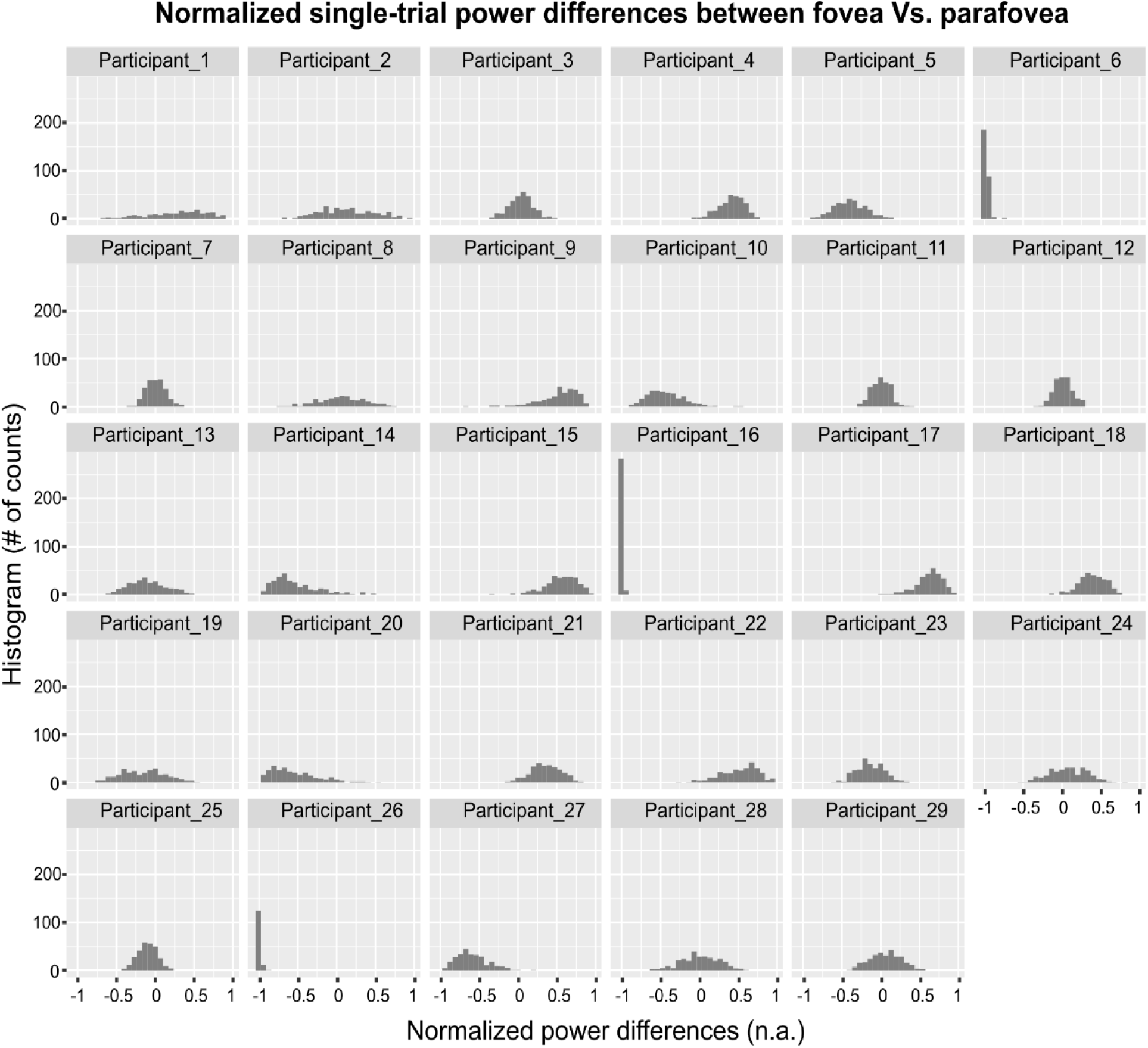
Histograms of the normalized single-trial power differences between the fovea and parafoveal, shown separately for each participant.

## Supplementary Notes

In Figure 4 of the Main Text, we observed significantly above-baseline tagging responses at both the foveal tagging frequency (*f_t_*) and the parafoveal tagging frequency (*f_p_*). Based on this finding, we proposed that attention was allocated in parallel to both the foveal and parafoveal words. However, tagging responses were measured using coherence, a cross-trial metric that quantifies the consistency between the flicker’s luminance changes (the tagging signal) and the MEG signal at a specific frequency. Because coherence reflects signal consistency across trials, it could produce high values for both *f_t_* and *f_p_* even if attention was allocated to the foveal word in some trials and to the parafoveal word in others. This leaves open the possibility that the observed coherence does not necessarily indicate simultaneous (parallel) attention allocation.

To address this concern, we turned to a single-trial measure: power at the tagging frequencies. For the subset of participants with RIFT (rapid invisible frequency tagging) response sensors at both *f_t_* and *f_p_* (*n* = 29), we calculated single-trial power at the foveal (𝑃𝑜𝑤_𝑡_) and parafoveal (𝑃𝑜𝑤_𝑝_) tagging frequency. Power values were computed by squaring the absolute values of the analytic signals obtained via Hilbert transformation, using the same procedure as in the coherence analysis. Then, for each participant, we averaged the single-trial power values across tagging response sensors and across the time window of 0 - 0.2s (aligned with fixation onset to target word). To compare these values, we calculated the normalised power difference for each trial: 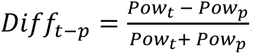. The normalisation step helps mitigate variability due to differences in signal-to-noise ratio across trials and participants. This index quantifies the relative similarity in power at the two tagging frequencies. The distribution of 𝐷𝑖𝑓𝑓_𝑡−𝑝_is shown in the grey panels in Supplementary figure 3 (distributions of individual participants in Supplementary figure 4). A unimodal distribution centred near zero would indicate similar power values and, by implication, similar amount of attention being allocated to both the fovea and parafovea. In contrast, a bimodal distribution, with one peak centred around a positive value and the other cantered around a negative value, would suggest trial-by-trial shifts of attention between the two regions.

In order to examine the statistical difference between power at these two frequencies, we fitted a linear mixed model using the *lmer* function from the lme4 package in R, with 𝐷𝑖𝑓𝑓_𝑡−𝑝_ as the dependent variable. The model included a random intercept for subject to account for repeated measurements within participants (𝑙𝑚𝑒𝑟(𝐷𝑖𝑓𝑓_𝑡−𝑝_ ∼ 1 + (1 | 𝑆𝑢𝑏𝑗𝑒𝑐𝑡𝑠)). The result did not reveal a significant deviation of 𝐷𝑖𝑓𝑓_𝑡−𝑝_ from zero (*b* = -0.065, *SE* = 0.086, *t*_(28)_ = -0.753, *p* = .458), suggesting that power at these two frequencies was comparable on a single-trial basis.

However, is the single-trial power measure reliable enough to detect tagging responses? To address this concern, we computed the single-trial power at a non-tagging frequency (55Hz), used as a noise estimation (𝑃𝑜𝑤_𝑛_), since *f_t_* and *f_p_* were either 60 or 65 Hz. We computed normalised single-trial power differences between foveal tagging and noise 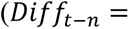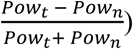 and between parafoveal tagging and noise 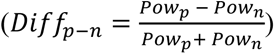. For distributions of these normalized power differences, see Supplementary figure 3. To assess whether 𝑃𝑜𝑤_𝑡_ and 𝑃𝑜𝑤_𝑝_were significantly above the noise level, we selected the lower of the two power values (foveal or parafoveal) on each trial and then compared it to the noise power at that trial. A linear mixed model (the same formula as above) revealed a significant effect (*b* = 3.126×10^-25^, *SE* = 3.958×10^-26^, *t*_(28)_ = 7.896, *p* = 1.350×10^-8^), indicating that both 𝑃𝑜𝑤_𝑡_ and 𝑃𝑜𝑤_𝑝_ were significantly higher than noise. This confirms that single-trial power estimation is a robust measure of RIFT responses.

In sum, this control analysis confirms that tagging responses at both fovea and parafovea were robust and comparable at the single-trial basis, providing further support for the notion that attention can be allocated to two words simultaneously.

